# Keratinocyte IL-17RA signaling promotes neutrophil recruitment and clinical lesions in experimental epidermolysis bullosa

**DOI:** 10.64898/2026.11.18.767503

**Authors:** Sandra Jukic, Alexandra Hölscher, Immo Prinz, Burkhard Becher, Ari Waisman, Enno Schmidt, Ralf J. Ludwig, Xinhua Yu, Frank Petersen, Christoph Hölscher, Fengyuan Deng

**Affiliations:** lnfection Immunology, Research Centre Borstel, Germany; Institute of Systems Immunology, Hamburg, University Medical Center, Hamburg-Eppendorf, Germany; Institute of Experimental Immunology, University of Zürich, Switzerland; Institute for Molecular Medicine, University Medical Center, Johannes Gutenberg University Mainz, Germany; Institute of Experimental Dermatology, University of Lübeck, Germany; Department of Dermatology, University Hospital Schleswig-Holstein, Campus Lübeck, Lübeck, Germany; Pulmonary Immune Diseases, Research Center Borstel, Borstel, Germany; Key Laboratory of Basic Research on Regional Diseases, Guangxi Medical University, Guangxi, China

**Author notes:** Correspondence to Christoph Hölscher (; Division of Infection Immunology, Research Center Borstel, Parkallee 33, 23845 Borstel, Germany) or Fengyuan Deng (; Key Laboratory of Basic Research on Regional Diseases, Guangxi Medical University, Guangxi, China).

**Keywords:** Epidermolysis bullosa acquisita, IL-17RA signaling, keratinocytes, neutrophil recruitment, antibody-transfer model

## Abstract

Epidermolysis bullosa acquisita (EBA) is an autoimmune blistering disease (AIBD) characterized by anti-type VII collagen autoantibody-mediated tissue injury. Neutrophils have been identified as essential for dermal-epidermal separation in various *in vitro* and mouse models of EBA. Although interleukin-17 (IL-17) signaling promotes neutrophilic inflammation, its contribution to EBA and the relevant responding cell types remain poorly defined. We investigated the role and cellular targets of IL-17 receptor A (IL-17RA) signaling using an antibody-transfer model of EBA in mice with global IL-17RA deficiency, combined IL-17A/IL-17F deficiency, IL-17F deficiency, or myeloid cell- or keratinocyte-specific IL-17RA deletion. Complementary *in vitro* assays assessed IL-17A-induced chemokine expression and neutrophil migration. Global IL-17RA deficiency, combined IL-17A/IL-17F deficiency, and IL-17F deficiency significantly reduced disease severity *in vivo*. Keratinocyte-specific IL-17RA deletion markedly attenuated EBA, whereas myeloid cell-specific deletion did not confer protection. *In vitro*, IL-17A increased *Cxcl1* mRNA expression in keratinocytes and fibroblasts, and conditioned media from stimulated cells promoted neutrophil migration. These findings identify keratinocytes as key cellular targets of IL-17RA signaling in experimental EBA and support a mechanism linking epithelial cytokine responses to neutrophil recruitment and tissue injury. Our data support investigating the keratinocyte IL-17RA pathway as a potential therapeutic target in EBA and related AIBD.

## INTRODUCTION

Epidermolysis bullosa acquisita (EBA) is a rare autoimmune blistering disease (AIBD) caused by autoantibodies against type VII collagen (COL7), a major structural component of anchoring fibrils at the dermal–epidermal junction (DEJ) (Chen et al., 2012; Kridin et al., 2019; Woodley et al., 1984). Clinically, EBA is characterized by skin fragility, tense blisters, erosions, and progressive scarring, which can cause substantial morbidity and impair quality of life (Koga et al., 2018; Vorobyev et al., 2017). The majority of patients present with the inflammatory variant of EBA resembling the bullous pemphigoid (BP), the by far most frequent AIBD. Disease pathogenesis involves an interplay between humoral autoimmunity and cellular inflammation, with neutrophils serving as key effector cells of tissue injury in mouse models of the disease mimicking the inflammatory variant of EBA (Hashimoto et al., 2012; Ludwig and Zillikens, 2011).

The interleukin-17 (IL-17) cytokine family comprises six members, IL-17A through IL-17F, that contribute to tissue inflammation and host defense (Chung et al., 2021; McGeachy et al., 2019). The best-characterized members, IL-17A and IL-17F, signal through a heterodimeric receptor complex consisting of IL-17 receptor A (IL-17RA) and IL-17 receptor C (IL-17RC) (Li et al., 2019; McGeachy et al., 2019). Ligand binding recruits the adaptor protein Act1 and activates downstream pathways involving NF-κB, mitogen-activated protein kinases (MAPKs), and C/EBP transcription factors (Gaffen, 2009). These pathways promote the expression of proinflammatory cytokines, chemokines, and antimicrobial peptides (Gaffen, 2009; Huangfu et al., 2023; Pappu et al., 2010).

IL-17 signaling has been implicated in several inflammatory and autoimmune diseases, including psoriasis, psoriatic arthritis, and bullous pemphigoid (Blauvelt and Chiricozzi, 2018; Brembilla et al., 2018; Chakievska et al., 2019; Heidari et al., 2025; Kong and Lai, 2024; Le Jan et al., 2014; Plee et al., 2015). Therapeutic targeting of IL-17 is established in plaque psoriasis and psoriatic arthritis, with approved agents such as the IL-17A inhibitors secukinumab and ixekizumab (Caso et al., 2025; Heidari et al., 2025). However, its contribution to pemphigoid disease such as EBA remains poorly understood. In particular, the cellular sources of pathogenic IL-17 and the cell types that respond to IL-17 signaling in EBA have not been clearly defined.

Neutrophil accumulation together with lymphocytes, eosinophils, and macrophages at the DEJ is a histopathological hallmark of EBA as an example of pemphigoid disease and is required for blister formation (Vorobyev et al., 2017). Neutrophil recruitment is regulated by CXC chemokines, particularly CXCL1 and CXCL2, released by resident skin cells in response to inflammatory stimuli (Allegretti, 2013). Epidermal keratinocytes act as immune sentinels and are important sources of these inflammatory mediators (Mercurio et al., 2020; Ye and Lai, 2025). Whether IL-17 signaling in keratinocytes is required to promote neutrophil recruitment and tissue injury in EBA remains unclear.

Here, we investigated the contribution of IL-17RA signaling to experimental EBA and identified the cell types through which it promotes disease. We combined global and cell type–specific genetic deletion models with in vitro functional assays to distinguish the roles of myeloid and keratinocyte IL-17RA signaling and to examine how IL-17A-induced responses in resident skin cells promote neutrophil migration.

## RESULTS

### Global IL-17RA deficiency attenuates experimental EBA

To determine the contribution of IL-17RA signaling to EBA, we induced antibody transfer-induced EBA in C57BL/6 and IL-17RA^-/-^ mice by subcutaneous administration of anti-mCOL7 IgG on days 0, 2, 4, 6, 8, and 10. Normal rabbit IgG served as a control. Following anti-mCOL7 IgG transfer, both strains developed progressive skin lesions, including hair loss, erythema, and crusting (Figure 1B). Disease severity was assessed as the percentage of affected body surface area on days 4, 8, and 12 as described (Kasprick et al., 2019). IL-17RA^-/-^ mice had significantly less affected skin than C57BL/6 mice. On day 12, mean clinical scores were approximately 55% lower in IL-17RA^-/-^ mice (C57BL/6: 7.8 ±3.5%; IL-17RA^-/-^: 3.4 ±1.2%; p < 0.05; Figure 1A). Histological analysis and neutrophil immunohistochemistry showed reduced neutrophil infiltration in lesional skin of IL-17RA^-/-^ mice, although perivascular infiltrates in both strains were predominantly neutrophilic (Figure 1B). Immunofluorescence (IF) staining detected tissue-bound IgG and C3 deposition at the DEJ in both anti-mCOL7 IgG-treated strains (Figure 1B). Lesional skin from IL-17RA^-/-^ mice also showed significantly reduced *Il6* and *Ccl2* mRNA expression compared with C57BL/6 mice (Figure 1C). The reduced mRNA expression documents changes associated with IL-17RA deficiency, but their functional contribution to disease severity was not assessed.

**Figure 1.**
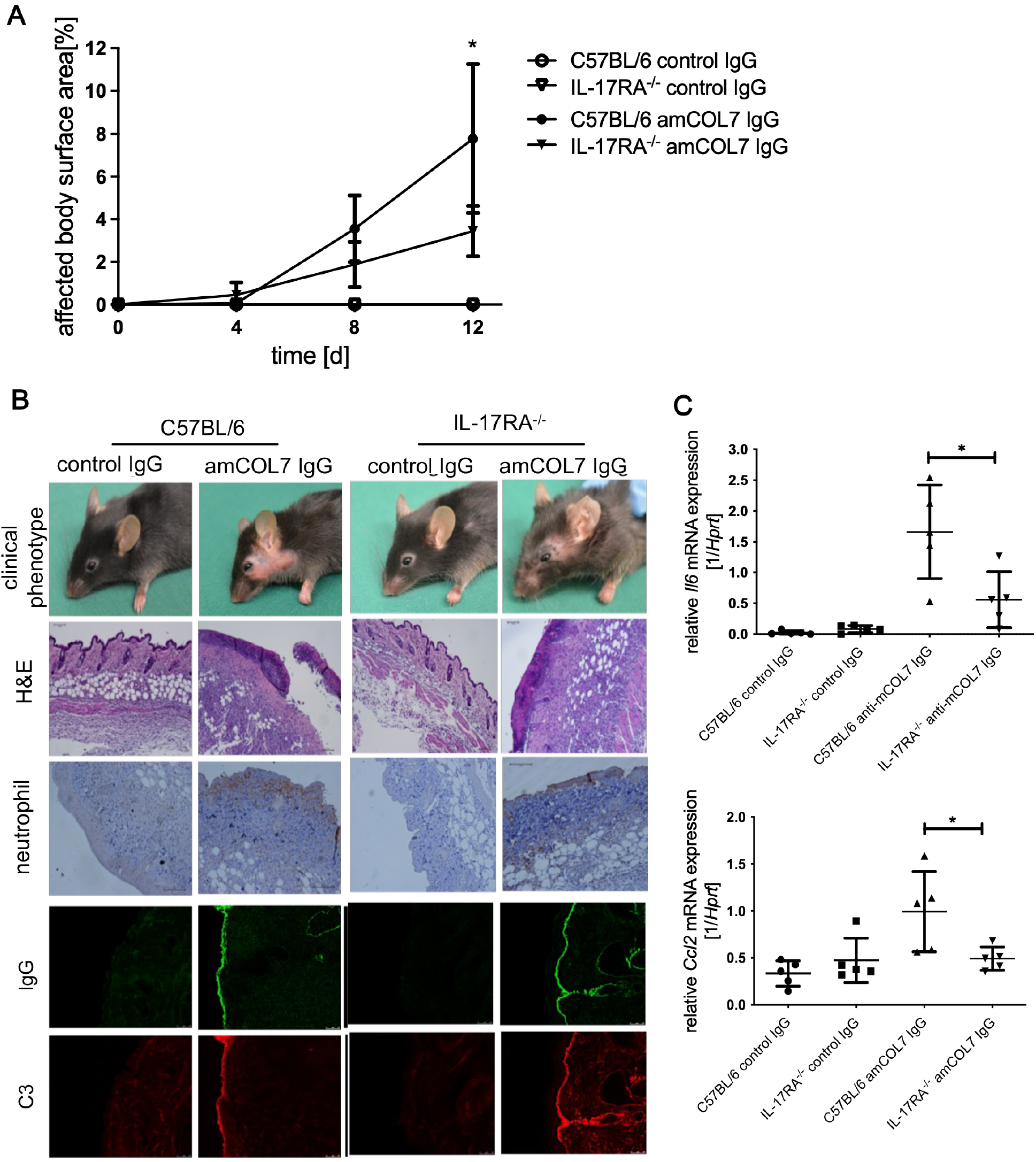
Global IL-17RA deficiency attenuates experimental EBA. C57BL/6 and global IL-17RA^−/−^ mice received six subcutaneous injections of anti-mCOL7 IgG (n = 5) or corresponding amounts of normal rabbit IgG (n = 5) every other day. (A) Clinical disease severity expressed as the percentage of affected body surface area over time. (B) Representative clinical photographs, hematoxylin and eosin (H&E)-stained ear skin sections, immunohistochemical staining of neutrophils in lesional skin, and immunofluorescence staining of tissue-bound IgG and complement C3 at the dermal–epidermal junction. Scale bar: 100 µm. (C) Relative mRNA expression of selected inflammatory cytokines and chemokines in lesional skin, measured by RT-qPCR. Data are presented as mean ± SD (n = 4–5 per group). *p < 0.05, **p < 0.01; Mann–Whitney rank-sum test.

### Both combined IL-17A/IL-17F deficiency or IL-17F deficiency reduce EBA severity

To examine the contribution of IL-17A and IL-17F to disease development, we induced antibody transfer-induced EBA in IL-17A/F^-/-^ mice. Similar to global IL-17RA deficiency, combined IL-17A/IL-17F deficiency significantly attenuated disease. On day 12, mean clinical scores were approximately 59% lower in double-deficient mice than in wild-type controls (C57BL/6: 14.5 ±3.9%; IL-17A/F^-/-^: 5.9 ±1.6%; p < 0.01; Figure 2A). Histological examination showed markedly reduced subepidermal blistering and neutrophil infiltration in IL-17A/F^-/-^ mice. Tissue-bound IgG and C3 deposition along the basement membrane zone (BMZ) were detected in both strains (Figure 2B). Lesional *Il6* and *Ccl2* mRNA expression was significantly lower in anti-mCOL7 IgG-treated IL-17A/F^-/-^ mice than in C57BL/6 controls. These changes were associated with combined IL-17A/IL-17F deficiency, but their functional contribution to disease severity was not assessed (Figure 2C).

**Figure 2.**
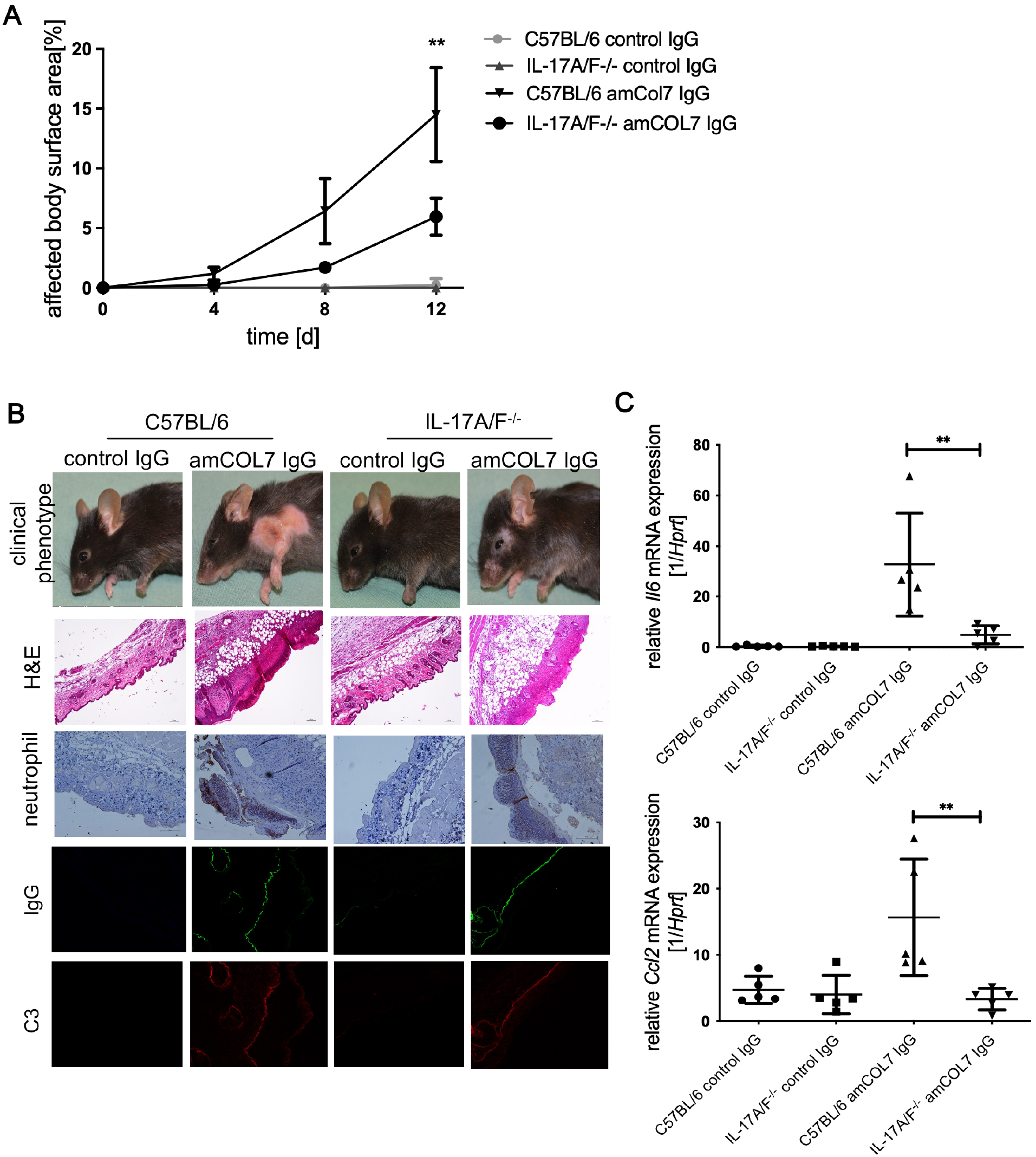
Combined IL-17A/IL-17F deficiency attenuates experimental EBA. C57BL/6 and IL-17A/F^−/−^ mice received six subcutaneous injections of anti-mCOL7 IgG (n = 5) or corresponding amounts of normal rabbit IgG (n = 5) every other day. (A) Clinical disease severity expressed as the percentage of affected body surface area over time. (B) Representative clinical photographs, hematoxylin and eosin (H&E)-stained ear skin sections, immunohistochemical staining of neutrophils in lesional skin, and immunofluorescence staining of tissue-bound IgG and complement C3 at the dermal–epidermal junction. Scale bar: 100 µm. (C) Relative mRNA expression of selected inflammatory cytokines and chemokines in lesional skin, measured by RT-qPCR. Data are presented as mean ± SD (n = 4–5 per group). *p < 0.05, **p < 0.01; Mann–Whitney rank-sum test.

Having previously demonstrated a pathogenic role for IL-17A in EBA (Deng et al., 2026), we next examined the contribution of IL-17F using IL-17F^-/-^ mice. Following anti-mCOL7 IgG transfer, IL-17F^-/-^ mice developed milder disease than C57BL/6 controls. The percentage of affected body surface area was significantly reduced on day 8 (C57BL/6: 2.5 ±1.2%; IL-17F^-/-^: 1.1 ±0.5%; p < 0.01) and day 12 (C57BL/6: 3.7 ±0.5%; IL-17F^-/-^: 2.2 ±1.0%; p < 0.01; Figure 3A). H&E staining showed inflammatory cell infiltration at the DEJ in both strains, and tissue-bound IgG and C3 deposition were assessed by IF microscopy (Figure 3B). Lesional *Ccl2* mRNA expression was markedly reduced in IL-17F^-/-^ mice compared with C57BL/6 controls. This change was associated with IL-17F deficiency, but its functional contribution to disease severity was not assessed (Figure 3C).

**Figure 3.**
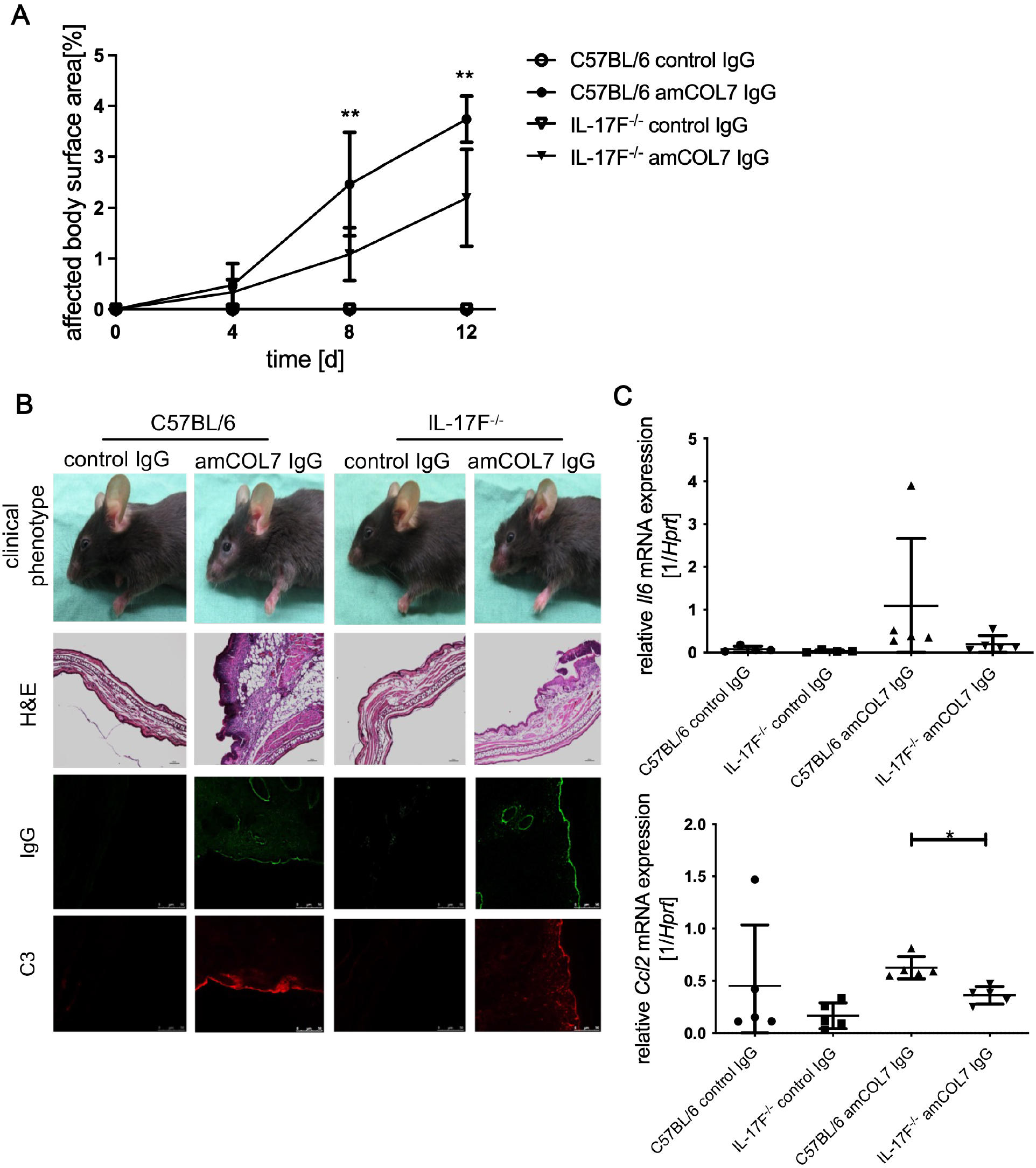
IL-17F deficiency attenuates experimental EBA. C57BL/6 and IL-17F^−/−^ mice received six subcutaneous injections of anti-mCOL7 IgG (n = 5) or corresponding amounts of normal rabbit IgG (n = 5) every other day. (A) Clinical disease severity expressed as the percentage of affected body surface area over time. (B) Representative clinical photographs, hematoxylin and eosin (H&E)-stained ear skin sections, and immunofluorescence staining of tissue-bound IgG and complement C3 at the dermal–epidermal junction. Scale bar: 100 µm. (C) Relative mRNA expression of selected inflammatory cytokines and chemokines in lesional skin, measured by RT-qPCR. Data are presented as mean ± SD (n = 4–5 per group). *p < 0.05, **p < 0.01; Mann–Whitney rank-sum test.

### Myeloid cell-specific IL-17RA deletion does not protect against EBA

Given the central role of neutrophils in experimental EBA, we investigated whether IL-17RA signaling in myeloid cells contributes to disease development. We crossed IL-17RA-floxed mice with LysM-Cre mice to generate myeloid cell-specific IL-17RA-deficient mice (IL-17RA^fl/fl^LysM-Cre^+^).

Following anti-mCOL7 IgG transfer, IL-17RA^fl/fl^LysM-Cre^+^ mice and Cre-negative littermate controls developed more severe disease than global IL-17RA^-/-^ mice. Clinical scores were significantly higher in myeloid cell-specific knockout mice than in global knockout mice (8.4 ±3.4% *vs* 4.1 ±2.3%, respectively; p < 0.05; Figure 4A). Myeloid cell-specific knockout mice and Cre-negative controls did not differ significantly in clinical phenotype, histological blister formation, neutrophil infiltration, or IgG and C3 deposition (Figure 4B,D). *Il6* and *Ccl2* mRNA expression did not differ among myeloid cell-specific knockout mice, Cre-negative littermates, and global IL-17RA^-/-^ mice (Figure 4C). Thus, myeloid cell-specific IL-17RA deletion did not confer protection against antibody transfer-induced EBA.

**Figure 4.**
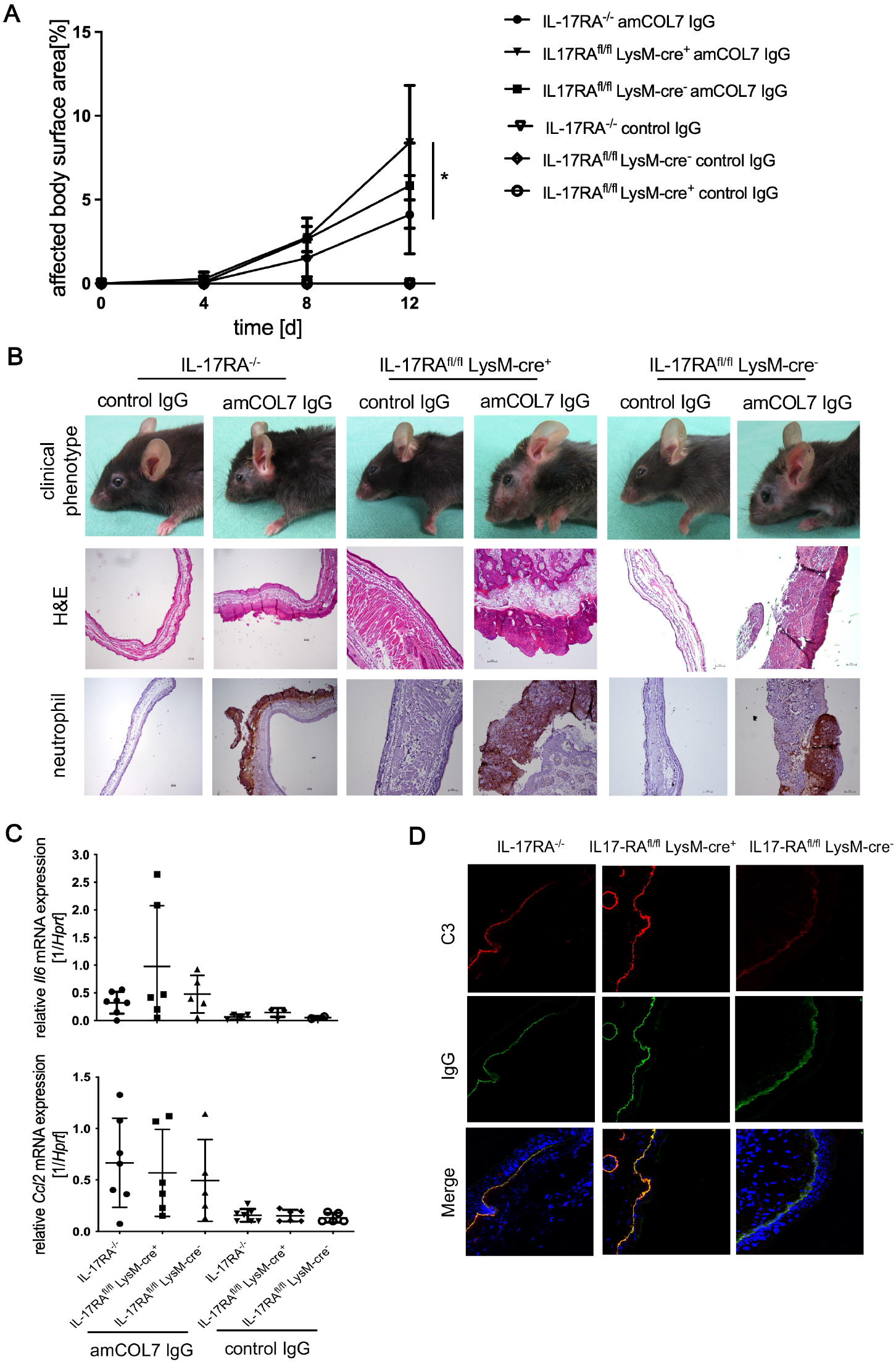
Myeloid cell-specific IL-17RA deletion does not protect against experimental EBA. IL-17RA^fl/fl^LysM-Cre^+^ mice, IL-17RA^fl/fl^LysM-Cre^−^ littermate controls, and global IL-17RA^−/−^ mice received six subcutaneous injections of anti-mCOL7 IgG (n = 5) or corresponding amounts of normal rabbit IgG (n = 5) every other day. (A) Clinical disease severity expressed as the percentage of affected body surface area over time. (B) Representative clinical photographs, hematoxylin and eosin (H&E)-stained ear skin sections, and immunohistochemical staining of neutrophils in lesional skin from all three genotypes. Scale bar: 100 µm. (C) Relative mRNA expression of selected inflammatory cytokines and chemokines in lesional skin, measured by RT-qPCR. (D) Immunofluorescence staining of tissue-bound IgG and complement C3 at the dermal–epidermal junction. Data are presented as mean ± SD (n = 4–5 per group). *p < 0.05, **p < 0.01; Mann–Whitney rank-sum test.

### Keratinocyte-specific IL-17RA deletion attenuates EBA

Because myeloid cell-specific IL-17RA deletion did not attenuate EBA, we next examined the contribution of IL-17RA signaling in resident skin cells. We crossed IL-17RA-floxed mice with K14-Cre mice to generate keratinocyte-specific IL-17RA-deficient mice (IL-17RA^fl/fl^K14-Cre^+^).

Keratinocyte-specific knockout mice developed markedly milder EBA than Cre-negative littermate controls. On day 12, clinical scores were significantly lower in IL-17RA^fl/fl^K14-Cre^+^ mice than in controls (4.4 ±2.2% vs 8.7 ±3.1%, respectively; p < 0.001; Figure 5A), with disease attenuation similar to that observed in global IL-17RA^-/-^ mice. Clinical and histological assessment showed reduced disease manifestations and minimal dermal–epidermal separation in keratinocyte-specific knockout mice (Figure 5B). *Il6* and *Ccl2* mRNA expression and tissue-bound IgG and C3 deposition did not differ significantly between groups (Figure 5C, D). These findings identify keratinocyte IL-17RA signaling as an important contributor to disease severity in experimental EBA.

**Figure 5.**
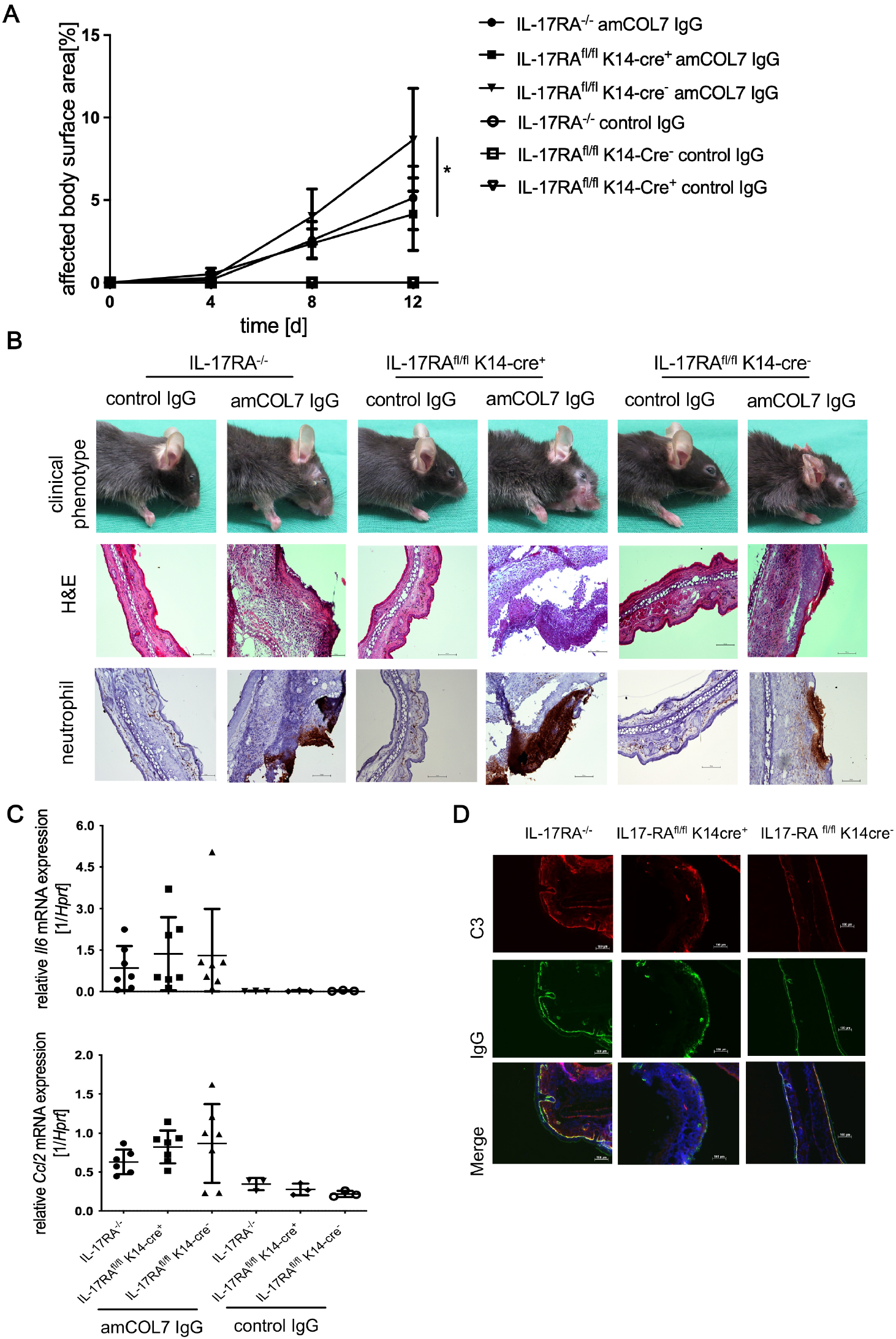
Keratinocyte-specific IL-17RA deletion attenuates experimental EBA. IL-17RA^fl/fl^K14-Cre^+^ mice, IL-17RA^fl/fl^K14-Cre^−^ littermate controls, and global IL-17RA^−/−^ mice received six subcutaneous injections of anti-mCOL7 IgG (n = 5) or corresponding amounts of normal rabbit IgG (n = 5) every other day. (A) Clinical disease severity expressed as the percentage of affected body surface area over time. (B) Representative clinical photographs, hematoxylin and eosin (H&E)-stained ear skin sections, and immunohistochemical staining of neutrophils in lesional skin from all three genotypes. Scale bar: 100 µm. (C) Relative mRNA expression of selected inflammatory cytokines and chemokines in lesional skin, measured by RT-qPCR. (D) Immunofluorescence staining of tissue-bound IgG and complement C3 at the dermal–epidermal junction. Data are presented as mean ± SD (n = 4–5 per group). *p < 0.05, **p < 0.01; Mann–Whitney rank-sum test.

### IL-17A increases chemokine mRNA expression in keratinocytes and fibroblasts

To investigate how IL-17RA signaling in resident skin cells may promote neutrophil recruitment, we assessed IL-17A-induced chemokine mRNA expression in murine keratinocytes and fibroblasts. Stimulation with recombinant IL-17A (100 ng/mL) for 24 hours significantly increased *Cxcl1* mRNA expression (Figure 6A, B; p < 0.001). These findings show that IL-17A induces a transcriptional response associated with neutrophil recruitment in resident skin cells.

**Figure 6.**
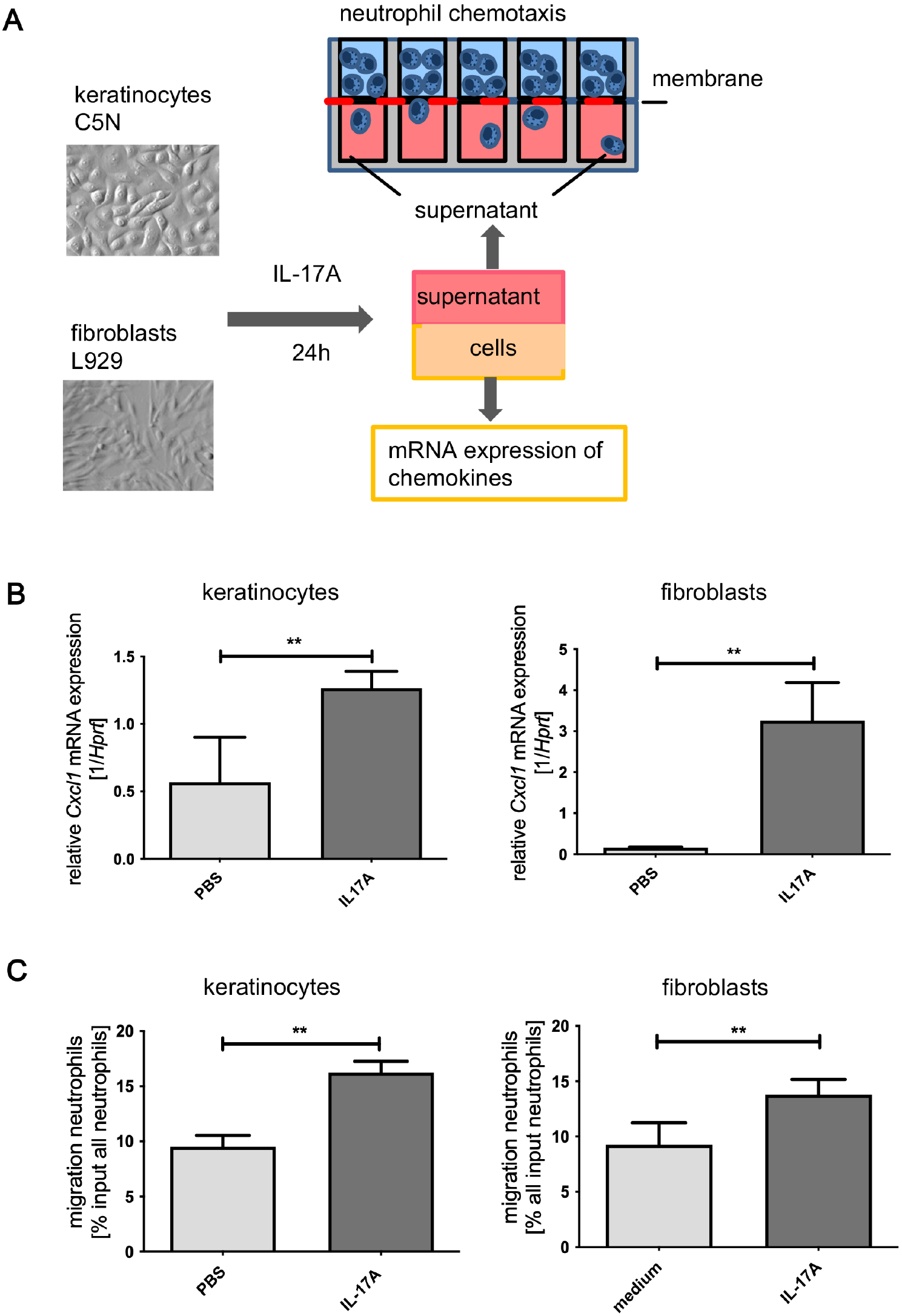
IL-17A increases chemokine mRNA expression in keratinocytes and fibroblasts, and conditioned media promote neutrophil migration in vitro. Keratinocytes and fibroblasts were cultured with or without IL-17A for 24 h. Cells were collected for mRNA analysis, and conditioned media were collected for neutrophil migration assays. (A) Schematic overview of the experimental design. Neutrophils were placed in the upper chamber and allowed to migrate through a membrane toward conditioned media in the lower chamber. (B) *Cxcl1* mRNA expression in keratinocytes and fibroblasts, measured by RT-qPCR. (C) Percentage of neutrophils migrating toward conditioned media from keratinocytes and fibroblasts after incubation for 1 h at 37 °C. Data are presented as mean ± SD. Each assay was performed in triplicate and repeated twice. **p < 0.01; t-test.

### Conditioned media from IL-17A-stimulated skin cells promote neutrophil migration

To assess whether IL-17A-induced responses in keratinocytes and fibroblasts promote neutrophil migration, we performed transwell assays using bone marrow-derived neutrophils. Conditioned media from IL-17A-stimulated keratinocytes and fibroblasts increased neutrophil migration approximately 1.5-to 2-fold compared with media from unstimulated cells (keratinocytes: 9.5 ±1.0% *vs* 16.2 ±1.0%, p < 0.01; fibroblasts, 9.3 ± 2.0% *vs* 13.8 ±1.4%, p < 0.01; Figure 6C). These findings demonstrate that conditioned media from IL-17A-stimulated resident skin cells have neutrophil-attracting activity, consistent with the observed increase in *Cxcl1* mRNA expression.

## DISCUSSION

In this study, we examined the cell type-specific contribution of IL-17RA signaling to EBA. Keratinocyte-specific IL-17RA deletion markedly attenuated clinical disease, whereas myeloid cell-specific deletion did not confer protection. Together with the reduced disease severity in globally IL-17RA-deficient mice, these findings identify keratinocytes as important cellular targets of pathogenic IL-17 signaling and support a mechanism linking epithelial responses to neutrophil recruitment and tissue injury.

The protection conferred by global IL-17RA deficiency and combined IL-17A/IL-17F deficiency is consistent with the established contribution of IL-17 signaling to neutrophil-mediated inflammation in experimental BP and psoriasis (Chakievska et al., 2019; van der Fits et al., 2009). These findings support a pathogenic role for IL-17RA signaling in antibody transfer-induced EBA as a model for pemphigoid disease. Reduced disease severity in IL-17F^-/-^ mice further indicates that IL-17F contributes to disease development. This observation is consistent with evidence that IL-17F induces epithelial chemokine production associated with neutrophil recruitment (Kawaguchi et al., 2003), although the mechanisms underlying its contribution to EBA were not directly examined here.

The conditional knockout experiments distinguish the contribution of myeloid IL-17RA signaling from the established effector functions of myeloid cells. Although IL-17 can promote myeloid cell recruitment and activation in inflammatory settings (Barin et al., 2012; McGinley et al., 2020), LysM-Cre-mediated IL-17RA deletion did not attenuate clinical or histopathological disease in our model. Thus, direct IL-17RA signaling in the targeted myeloid populations does not appear to be required for full disease expression under these experimental conditions. This finding does not imply that neutrophils or macrophages are dispensable for EBA; rather, it suggests that their pathogenic activity can be sustained without cell-intrinsic IL-17RA signaling. In patients with BP, neutrophils and T cells have been identified as cellular sources of IL-17A in lesional skin (Chakievska et al., 2019; Le Jan et al., 2014).

In contrast, the marked disease attenuation in keratinocyte-specific IL-17RA-deficient mice identifies an important contribution of epidermal cytokine sensing to EBA. Rather than serving only as targets of inflammatory injury, keratinocytes can actively amplify cutaneous inflammation through IL-17-induced chemokine responses (Cui et al., 2024). This interpretation aligns with studies highlighting their immunoregulatory functions in inflammatory skin disease (Cui et al., 2024; Kamata and Tada, 2023) and with evidence that IgG directed against COL17, the major autoantigen in BP, induces dose- and time-dependent release of IL-6 and IL-8 from cultured keratinocytes (Schmidt et al., 2000). Our findings extend this concept to experimental EBA and implicate keratinocyte IL-17RA signaling in the propagation of antibody-mediated tissue injury.

The distinct effects of keratinocyte- and myeloid cell-specific IL-17RA deletion identify a cell type-dependent contribution to experimental EBA, but the molecular basis of this distinction remains unresolved. IL-17A and IL-17F signal through the IL-17RA/IL-17RC complex, whose component availability can influence cellular responsiveness (Furue et al., 2020; Ho et al., 2010). Downstream signaling involves Act1 and TRAF6, with activation of NF-κB and mitogen-activated protein kinase pathways (Gaffen, 2009; Song et al., 2016). IL-17 can also prolong chemokine mRNA stability through mechanisms involving TRAF proteins, SF2/ASF, and HuR (Herjan et al., 2013; Sun et al., 2011), while negative regulators such as USP25 constrain signaling (Gaffen, 2009; Zhong et al., 2012). These mechanisms offer possible explanations for sustained epithelial chemokine responses; however, receptor abundance, signaling activity, and mRNA stability were not compared between keratinocytes and myeloid cells in this study. Myeloid cells can respond to other inflammatory cues, including immune-complex engagement of Fcγ receptors (Ludwig, 2013), and direct neutrophil IL-17 responsiveness has been described in other settings (Taylor et al., 2014). Thus, our findings support a prominent role for keratinocyte IL-17RA signaling in this model, not a universal lack of myeloid responsiveness to IL-17.

Our *in vitro* assays complement the *in vivo* findings: IL-17A increased *Cxcl1* mRNA expression in keratinocytes and fibroblasts, and conditioned media from stimulated cells enhanced neutrophil migration. These results are consistent with the generation of neutrophil-attracting signals, although they do not establish CXCL1 as the specific mediator of migration. Once recruited to the cutaneous BMZ, neutrophils can be activated by local immune complexes (Kasperkiewicz et al., 2016; Ludwig, 2013). Engagement of Fcγ receptors by COL7-bound autoantibodies promotes degranulation and an oxidative burst (Haeger et al., 2022; Hashimoto et al., 2012). Reactive oxygen species and granule-associated mediators, including neutrophil elastase, matrix metalloproteinase-9, and myeloperoxidase, contribute to inflammatory injury (Hashimoto et al., 2012; Kasperkiewicz et al., 2016; Shimanovich et al., 2004). Proteolytic disruption of BMZ components compromises DEJ integrity (Hashimoto et al., 2012; Taylor et al., 2014), while neutrophil extracellular traps may further amplify local inflammation (Domínguez-Díaz et al., 2021). Together, these observations support a model in which keratinocyte IL-17RA signaling promotes recruitment, followed by immune-complex-mediated neutrophil activation and tissue injury, rather than establishing this pathway as the sole requirement for neutrophil accumulation.

These findings provide a rationale for investigating the therapeutic potential of disrupting the keratinocyte–neutrophil axis in EBA. Biologics targeting IL-17A, including secukinumab and ixekizumab, or IL-17RA, such as brodalumab, are effective in psoriasis and have been investigated in other inflammatory skin diseases including BP (Brembilla and Boehncke, 2023; Heidari et al., 2025; Holtsche et al., 2026; Kasperkiewicz et al., 2016; Papp et al., 2012). However, genetic protection in our model does not establish the efficacy or safety of these agents in EBA, which requires direct therapeutic evaluation and validation in human disease. The contribution of keratinocyte IL-17RA signaling also raises the possibility of topical or epithelium-targeted inhibition of downstream pathways, including Act1-dependent signaling. Whether such approaches can suppress local chemokine responses while limiting systemic effects remains to be determined.

Several limitations should be acknowledged. The antibody-transfer model primarily captures the neutrophil-driven effector phase of EBA, whereas lesional skin in human EBA also contains lymphocytes, eosinophils, and macrophages. The relevance of our findings to human disease therefore remains to be established. Although we focused on IL-17A and IL-17F, other IL-17RA-dependent ligands may contribute. In particular, the epithelial-derived cytokine IL-17C signals through IL-17RE/IL-17RA and warrants investigation (Chang et al., 2011). The contribution of other resident skin cells, including dermal fibroblasts and endothelial cells, to local CXC chemokine responses also remains to be quantified. Finally, the *in vitro* findings establish increased *Cxcl1* mRNA expression and enhanced neutrophil migration toward conditioned media, but not CXCL1 protein secretion or a causal requirement for CXCL1 in migration. The individual factors responsible for the migratory response therefore remain undefined.

In conclusion, our study identifies keratinocyte IL-17RA signaling as an important contributor to inflammatory tissue injury in experimental EBA. The contrasting effects of keratinocyte- and myeloid cell-specific deletion, together with the activity of conditioned media from IL-17A-stimulated skin cells, are consistent with an indirect mechanism of neutrophil recruitment through resident skin cells. However, these experiments do not establish the complete causal pathway linking keratinocyte IL-17RA signaling, individual chemokines, and neutrophil recruitment in vivo. These findings refine our understanding of the cellular basis of antibody-mediated skin inflammation and highlight the keratinocyte IL-17RA pathway as a potential therapeutic target requiring further preclinical and clinical validation.

## MATERIALS AND METHODS

### Mice

*Il17ra*^tm1.2Kurs^ (IL-17RA^−/−^) (El Malki et al., 2013), *Il17a/Il17f*^tm1.1Impr^ (IL-17A/F^−/−^) (Haas et al., 2012) and *Il17f*^tm1Awai^ (IL-17F^−/−^) (Haak et al., 2009) mice were bred at the animal facility of the Christian-Albrechts-University Kiel (Kiel, Germany). For these mice, C57BL/6 mice served as controls. *Il17ra*^tm1.1Kurs^ (IL-17RA^fl/fl^) (El Malki et al., 2013), *Lyz*^tm1Ifo^ (LysM-Cre) (Clausen et al., 1999), and Tg(KRT14-cre)1Amc (K14-Cre) mice (Dassule et al., 2000) were bred at the animal facility of the Research Center Borstel (Borstel, Germany). To generate macrophage/neutrophil-specific and keratinocyte-specific IL-17RA-deficient mice, IL-17RA^fl/fl^ were crossed to LysM-Cre or K14-Cre mice, respectively. Cre-negative littermates were used as control mice. All mice used in the experiments were female and 8–16 weeks old. All experimental animals were housed under specific pathogen-free conditions with a 12-hour light/dark cycle at the animal facility of the Research Center Borstel. All animal experiments were reviewed and approved by the Animal Research Ethics Board of the Ministry of Agriculture, Rural Areas, European Affairs and Consumer Protection, Kiel, Germany (approval numbers 12-2/14 and 15-2/18). Clinical examinations, biopsies, and blood sampling were performed under anesthesia induced by intraperitoneal administration of ketamine (100 μg/g) and xylazine (15 μg/g).

### Anti-mCOL7 antibodies

Rabbit anti-mCOL7c antibodies and control rabbit IgG were prepared as described previously (Sitaru et al., 2005). Briefly, New Zealand White rabbits were immunized subcutaneously with recombinant mCOL7c protein (amino acid residues 757–967) emulsified in Freund’s complete adjuvant, followed by two booster immunizations with mCOL7c protein emulsified in Freund’s incomplete adjuvant. Total IgG was purified from sera of immunized rabbits by affinity chromatography using Protein G Sepharose Fast Flow. Control rabbit IgG was isolated from sera of non-immunized rabbits. The reactivity of all IgG fractions was assessed by immunofluorescence microscopy using murine skin.

### Induction and evaluation of experimental EBA

Experimental EBA was induced as described previously (Kasprick et al., 2019), with minor modifications. Female mice aged 8–16 weeks received six subcutaneous injections of rabbit anti-mCOL7 IgG (2–3 mg per injection, with exact doses specified in the respective experimental protocols) or normal rabbit IgG into the abdomen on days 0, 2, 4, 6, 8, and 10. Disease severity was assessed on days 4, 8, and 12 and expressed as the percentage of affected body surface area. Cutaneous lesions, crusting, erythema, and alopecia were evaluated using the following body-region weights: 2.5% each for the ears, snout, and oral mucosa; 0.5% for each eye; 9% for the head and neck, excluding the eyes, ears, snout, and oral mucosa; 5% for each forelimb; 10% for the hind limbs and tail; and 40% for the trunk (Gross et al., 2024). The regional scores were summed to obtain the overall clinical score. On day 12, mice were sacrificed, and sera and biopsies were collected.

### Hematoxylin and eosin staining of lesional skin

Skin and ear samples were fixed in 4% buffered formaldehyde and embedded in paraffin. Sections (5 μm thick) were stained with hematoxylin and eosin (H&E) according to standard protocols and evaluated histologically.

### Immunohistochemical detection of neutrophil infiltration

Formalin-fixed, paraffin-embedded sections of lesional skin were deparaffinized in xylene and rehydrated through graded ethanol. Antigen retrieval was performed in 10 mM citrate buffer (pH 6.0) at 121 °C for 1 h. Endogenous peroxidase activity, endogenous biotin, and nonspecific binding were blocked with 3% H_2_O_2_, biotin blocking solution (Vector Laboratories), and 5% BSA, respectively. Sections were incubated overnight at 4 °C with an anti-murine neutrophil antibody (07-APR, Cedarlane), followed by a biotinylated secondary antibody (Jackson ImmunoResearch) for 45 min at room temperature and avidin-biotinylated HRP (Vector Laboratories) for 20 min. Staining was visualized with DAB (Vector Laboratories). Sections were counterstained with hematoxylin for 5 min, and images were acquired by bright-field microscopy (Nikon).

### Detection of tissue-bound IgG and complement

Tissue-bound rabbit IgG and complement C3 deposition were assessed by immunofluorescence staining of murine skin cryosections (5 µm thick). After blocking with Roti-Immunoblock (Roth), rabbit IgG was detected using Alexa Fluor 488-conjugated goat anti-rabbit IgG (Invitrogen). Complement C3 was detected using a rat anti-mouse C3 antibody (Cedarlane), followed by Alexa Fluor 546-conjugated goat anti-rat IgG (Invitrogen). Sections were mounted with Gold mounting medium (Thermo Fisher), and fluorescence images were acquired using a Leica SP5 confocal microscope.

### Quantification of mRNA expression in skin

Gene expression in lesional skin was assessed by RNA isolation, reverse transcription, and quantitative real-time PCR (RT-qPCR) as described previously (Erdmann et al., 2018). Total RNA was isolated from skin biopsies using the RNeasy Mini Kit (Qiagen) according to the manufacturer’s instructions. Following reverse transcription, cDNA was amplified using a LightCycler 480 system (Roche). Each reaction contained 1 µL of cDNA template or standard and 9 µL of master mix containing 50% 2× LC480 Probes Master, 6.25 µM sense and antisense primers, and 1% Universal ProbeLibrary probe in ddH2O. The following primer and probe sets were used. *Hprt*: sense 5′-TCC TCC TCA GAC CGC TTT T-3′, antisense 5′-CCT GGT TCA TCA TCG CTA ATC-3′, probe #95; *Il1b*: sense 5′ -TGT-AAT-GAA-AGA-CGG-CAC-ACC-3′, antisense 5′-TCT-TCT-TTG-GGT-ATT-GCT-TGG-3 ′ , probe #78; *Il6*: sense 5 ′ -GTC ACC AAA CTG GAT ATA ATC AGG A-3 ′ , antisense 5′-CCA GGT AGC TAT GGT ACT CCA GAA-3′, probe #6; *Cxcl1*: sense 5′-AGA-CTC-CAG-CCA-CAC-TCC-AA-3′, antisense 5′-TGA-CAG-CGC-AGC-TCA-TTG-3′, probe #83; *Cxcl2*: sense 5′-CCA-GCC-ACA-CTT-CAG-CCT-A-3′, antisense 5′-CAG-TTC-ACT-GGC-CAC-AAC-AG-3′, probe #49; *Ccl2*: sense 5′-CAT-CCA-CGT-GTT-GGC-TCA-3′, antisense 5′-GAT-CAT-CTT-GCT-GGT-GAA-TGA-GT-3′, probe #62. Data were analyzed using LightCycler software LCS480 version 1.5.0.39 (Roche).

### Fibroblast and keratinocyte assay

Murine keratinocytes (C5N) and murine fibroblasts (L929) were cultured in 48-well plates at a density of 10 × 10^6^ cells/mL and stimulated with IL-17A (100 ng/mL). PBS served as the control. Cells were collected for RT-qPCR analysis of Cxcl1 and Cxcl2 mRNA expression. Culture supernatants were collected 24 h after stimulation and used in the neutrophil migration assay.

### Neutrophil transwell migration assay

Bone marrow-derived neutrophils from wild-type mice were resuspended at 4 × 10^6^ cells/mL in complete medium supplemented with G-CSF (10 ng/mL) and incubated at 37 °C in 5% CO_2_ for 48 h. Cells were then collected using a discontinuous Percoll gradient. Neutrophil migration was assessed using a 48-well macrochemotaxis chamber (Neuro Probe) with nitrocellulose filters (Neuro Probe). The chemotactic solution was diluted in PBS and added to the lower wells (30 µL per well). PVP-treated nitrocellulose filters with a pore size of 3 µm were placed between the lower and upper plates. Neutrophils were added to the upper wells (30 µL per well) at 2 × 10^6^ cells/mL. The chamber was incubated at 37 °C in 5% CO_2_ for 1h. Migrated cells in the lower wells were counted, and the percentage of migrated neutrophils was calculated.

### Statistical analysis

Statistical analyses were performed using GraphPad Prism version 6.0 (GraphPad Software). Unless otherwise stated, data are presented as mean ± standard deviation (SD). Two-group comparisons were performed using Student’s t-test or the Mann–Whitney rank-sum test, as appropriate. For comparisons among more than two groups, normally distributed data were analyzed by one-way ANOVA followed by Bonferroni’s multiple-comparison test. Nonparametric data were analyzed using the Kruskal–Wallis test followed by Bonferroni’s multiple-comparison test. Statistical significance was indicated as follows: *p < 0.05, **p < 0.01, and ***p < 0.001.

## ETHICS STATEMANT

All animal experiments were reviewed and approved by the Animal Research Ethics Board of the Ministry of Agriculture, Rural Areas, European Affairs and Consumer Protection of the State of Schleswig-Holstein, Kiel, Germany (approval numbers 91-9/10, 4-1/13, 12-2/14, and 109-10/19).

## DATA AVAILABILITY

The data supporting the findings of this study are available from the corresponding author upon reasonable request.

## CONFLICTS OF INTEREST

All authors declare that they have no conflicts of interest.

## ACKNOWLEDGMENTS

We are grateful to Anja Walter and Marion Schuldt for supplying and cleaning the lab and to Ika Monath and Christine Keller for organizing the animal facility at the Research Center Borstel. This work was supported by the GRK 1727 "Modulation of Autoimmunity" (Project-ID 179309734).

## AUTHO CONTRIBUTIONS

Conceptualization: CH, XY, and FP; Supervision: CH; Investigation: FD, SJ, AH; Formal analysis: FD, SJ; Resources: IP, BB, AW, RL, and ES; Visualization: FD; Writing: FD; Writing - review & editing: CH, XY, FP, IP, AW, RL and ES.

## DECLARATION OF GENERATIVE ARTIFICIAL INTELLIGENCE (AI) OR LARGE LANGUAGE MODELS (LLMS)

During the preparation of this manuscript, the authors used Microsoft Copilot to assist with language editing and improving readability. All AI-assisted content was critically reviewed and edited by the authors, who take full responsibility for the content of the publication.

## Notes

### Competing Interest Statement

The authors have declared no competing interest.

